# KP.2-based monovalent mRNA vaccines robustly boost antibody responses to SARS-CoV-2

**DOI:** 10.1101/2024.12.02.626472

**Authors:** Qian Wang, Ian A. Mellis, Madeline Wu, Anthony Bowen, Carmen Gherasim, Riccardo Valdez, Jayesh G. Shah, Lawrence J. Purpura, Michael T. Yin, Aubree Gordon, Yicheng Guo, David D. Ho

## Abstract

In response to the ongoing evolution of SARS-CoV-2, COVID-19 mRNA vaccines were recently updated to encode the spike protein of the KP.2 subvariant of the JN.1 sublineage. However, the immunogenicity of KP.2-based monovalent mRNA vaccines (KP.2 MV) has yet to be fully evaluated and reported, particularly against dominant and growing viral variants KP.3.1.1 and XEC, which bear some distinct mutations from KP.2. Here we report that KP.2 MV boosters elicit robust neutralizing antibody titers in a cohort of 16 healthy adult participants against all tested variants in pseudovirus neutralization assays. The highest post-boost geometric mean titers were against older variants D614G (17,293) and BA.5 (14,358), suggestive of immune imprinting, but the post-boost titers against currently dominant or growing viruses KP.3.1.1 (1,698) and XEC (1,721) were still robust. Fold-changes in titers were highest against recent JN.1 subvariants, including JN.1, KP.2, KP.3, KP.3.1.1, and XEC, (5.8-to-7.8-fold), compared to older variants D614G and BA.5 (1.6- and 2.5-fold), which suggests that KP.2 MV boosters have at least partially mitigated immune imprinting. Overall, these results show that KP.2 MV boosters elicit robust neutralizing antibodies against dominant SARS-CoV-2 viruses.

## Main Text

In response to the ongoing evolution of SARS-CoV-2, vaccine manufacturers have released updated COVID-19 vaccines annually since 2022. For much of 2024, the global spread has been dominated by the JN.1 lineage of viruses^1^, which are antigenically quite distant from the XBB.1.5 variant that was used in the previous vaccine booster^2^. In August 2024, the U.S. Food and Drug Administration authorized two updated mRNA vaccines (Pfizer-BioNTech and Moderna) based on the spike sequence of KP.2, a subvariant in the JN.1 lineage^3^. In the United Kingdom and the European Union, a KP.2-based mRNA vaccine (BioNTech) was also authorized later in the year^4,5^. We now provide the first indication of the acute boosting effect of updated KP.2 monovalent mRNA vaccines (KP.2 MV) on serum SARS-CoV-2 neutralizing antibodies in humans.

Since the authorization of the updated vaccine boosters, SARS-CoV-2 has evolved beyond KP.2, with subvariant KP.3.1.1 becoming dominant globally and the subvariant XEC now gaining traction rapidly^1^. KP.2 contains R346T, F456L, and V1104L mutations in spike in addition to those present in the parental JN.1 (**Figure 1A**). Both KP.3.1.1 and XEC share F456L and V1104L found in KP.2, along with Q493E, which is absent in KP.2. In addition, KP.3.1.1 harbors the S31 deletion (S31Δ), while XEC carries T22N and F59S; neither KP.3.1.1 nor XEC possesses the R346T mutation (**Figure 1A**). The effectiveness of the updated KP.2 MV boosters on neutralizing antibodies in human sera against recently dominant subvariants has yet to be reported.

**Figure 1.**
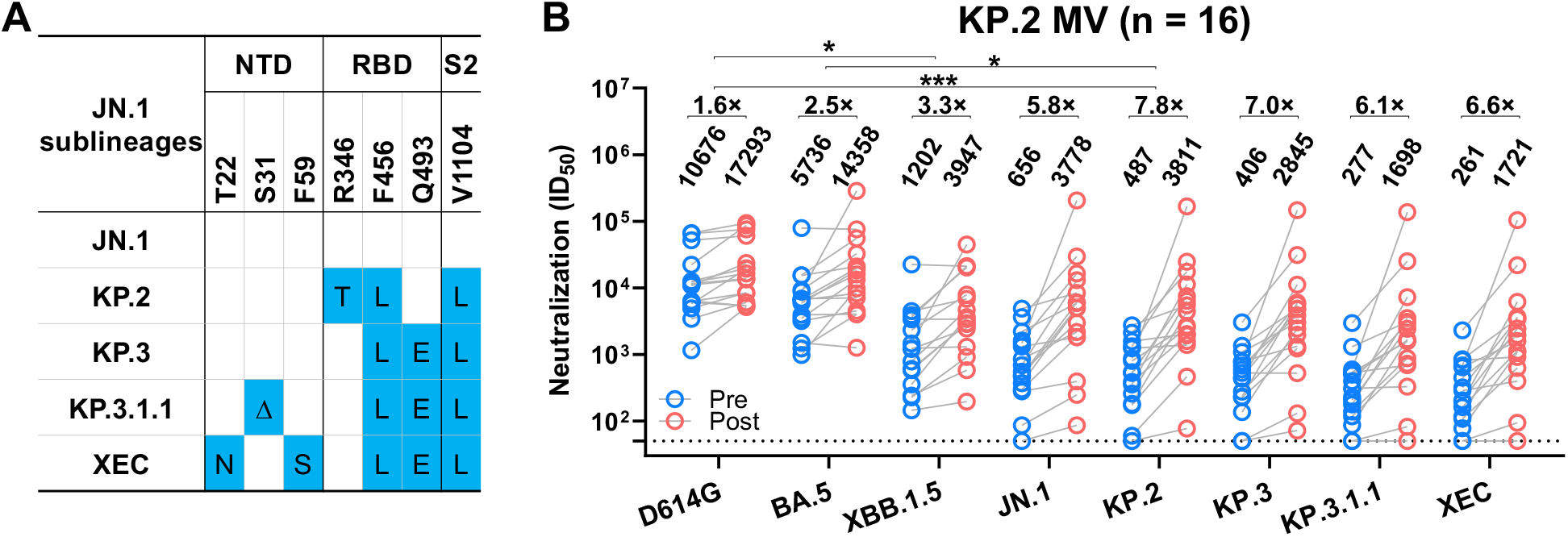
SARS-CoV-2 neutralizing antibody responses before and after a KP.2 monovalent mRNA vaccine booster. **A.** Spike mutations of the indicated JN.1 sublineages. NTD, N-terminal domain; RBD, receptor-binding domain; Δ, deletion. **B.** Serum virus-neutralizing titers (ID_50_) against the indicated SARS-CoV-2 pseudoviruses and the fold changes between pre- and post-vaccination serum samples are shown. Comparisons of fold increases in ID_50_ titers after KP.2 MV between KP.2 and other variants were analyzed using the Mann-Whitney U tests. *p < 0.05; ***p < 0.001; P > 0.05 are not shown. MV, monovalent vaccine. n, sample size. The dotted line represents the assay limit of detection (LOD) of 50.

To assess serum neutralizing antibodies elicited by KP.2 MV boosters, vesicular stomatitis virus (VSV)-pseudotyped viruses were generated for JN.1, KP.2, KP.3, KP.3.1.1, and XEC, as well as reference viruses D614G, BA.5, and XBB.1.5^1,6^. These pseudoviruses were subjected to neutralization assays using serum samples collected from 16 healthy volunteers before vaccination, as well as about a month (mean 30.9 days; range 26-39 days) post KP.2 MV booster (**Tables S1 and S2**). Vaccination boosted virus-neutralizing 50% inhibitory dilution (ID_50_) titers in serum by 1.6-fold against D614G, 2.5-fold against BA.5, and 3.3-fold against XBB.1.5 (**Figure 1B**), showing a persistent but modest back-boosting effect. Notably, a more substantial boost (6.1-to 7.8-fold) was observed for subvariants in the JN.1 lineage: KP.2, KP.3, KP.3.1.1, and XEC, a finding that suggests the new boosters have seemingly mitigated the immunological imprinting problem observed with previous bivalent COVID-19 vaccine boosters^6^.

No significant differences in boost effects by KP.2 MV were observed between age groups (<40 years vs. >40 years), self-reported genders (female vs. male), or vaccine manufacturers (Moderna vs. Pfizer-BioNTech) (**Figure S1**).

Overall, serum neutralizing titers against KP.3.1.1 and XEC were similarly lower than titers against JN.1, KP.2, and KP.3 in both pre and post samples (**Figure 1B**). This finding of greater antibody evasion is likely one explanation for their recent dominance in the population. Post boost, the geometric mean serum ID_50_ titers against KP.3.1.1 and XEC were quite robust at 1,698 and 1,712, respectively. Such levels of neutralizing antibodies are predictive of protection against symptomatic COVID-19 infection in prior clinical studies or vaccine trials trials^6-8^, although the exact levels that correlate with protection do vary according to the neutralization assays employed^6-8^.

In summary, a KP.2 MV booster elicits robust short-term neutralizing antibodies against recent SARS-CoV-2 subvariants KP.3.1.1 and XEC variants. The durability of this booster response needs to be assessed in time, as well as the associated clinical efficacy. The relentless evolution of SARS-CoV-2 underscores the challenge of continually updating our vaccines for optimal protection.

