## Supplementary Appendix for "KP.2-based monovalent mRNA vaccines robustly boost antibody responses to SARS-CoV-2"

### Supplemental Appendix

#### Supplemental Methods

*Clinical cohorts*

Serum samples were collected as part of the VIVA at the University of Michigan and as part of the "COVID-19 Persistence and Immunology Cohort (C-PIC)" study at Columbia University. Specimens were obtained following participant consent and in adherence to the protocols approved by the IRBs of University of Michigan Medical School (protocol HUM00232359) and Columbia University (protocol AAAS9722).

In this study, serum samples were collected from individuals who had been administered the KP.2 monovalent vaccine booster (KP.2 MV). Serum was collected both before and after receiving the booster. The majority of the study subjects were female, representing 56.3%, with an average age of 41.4 years. Serum samples were collected, on average, 14.4 days pre KP.2 MV booster and 30.9 days post KP.2 MV booster. Further demographic details, vaccination status, and serum collected timelines can be found summarized in the **Table S1 and S2**.

*Cell lines*

HEK293T (ATCC, CRL-3216) cells and Vero-E6 cells (ATCC, CRL-1586) were cultured in Dulbecco's Modified Eagle's Medium (DMEM) that contained 10% heat-inactivated fetal bovine serum (FBS) and 1% penicillin-streptomycin (PS). All cell lines were cultured in an atmosphere of 5% $CO_2$ at 37°C.

*SARS-CoV-2 spike plasmids*

The spike constructs of D614G, BA.5, XBB.1.5, JN.1, KP.2, KP.3, KP.3.1.1 and XEC were generated as previously reported[1-3]. All constructs were confirmed by Sanger sequencing.

*Pseudotyped SARS-CoV-2 variants*

Pseudotyped SARS-CoV-2 was produced following a previously established protocol[4]. HEK293T cells were first transfected with spike-encoding plasmids using 1 mg/mL of PEI-MAX (Polysciences, Inc.) and cultured for 24 hours. The transfected HEK293T cells were then infected with VSV-G pseudotyped ΔG-luciferase virus (Kerafast, EH1020-PM) at a multiplicity of infection (MOI) of approximately 3 to 5. Two hours later, the cells were washed three times with and cultured in fresh competent cell medium overnight. The virus was then harvested first by centrifugation at 2000 rpm for 10 minutes followed by collection of the supernatant. 20% of I1 hybridoma (ATCC, CRL-2700) supernatant was then added to each virus before storing at -80°C until use.

*Pseudovirus neutralization*

Pseudoviruses were titrated to standardize the viral input prior to each neutralization assay. For neutralization assays, serum samples were inactivated at 56°C for 30 minutes before use, and the inactivated sera were serially diluted with a dilution factor of four on black Co-Star neutralization plates. Following this, pseudoviruses were added and incubated at 37 °C for 1 hour. As a control, wells containing only the pseudovirus were also prepared on each test plate. Vero-E6 cells were then seeded at 40,000 cells per well and were incubated overnight at 37°C. Afterwards, cellular lysis was conducted, and the resultant luciferase activity was quantified employing the Tecan Infinite® 200 PRO using i-control™ software v.3.9.1.0, in accordance with the manufacturer's instructions. The serum dilution that inhibits 50% of virus entry ($ID_{50}$) was calculated using a five-parameter dose-response curve fitting using GraphPad Prism v.10.3.

*Quantification and statistical analysis*

$ID_{50}$ values at or below the assay limit of detection (LOD) of 50 were treated as 50 for the purpose of calculating each group's geometric mean $ID_{50}$ titer (GMT). Statistical significance between was evaluated using the Mann-Whitney unpaired t tests in GraphPad Prism v.10.3. Levels of significance are denoted as follows: ns, not significant; $*p < 0.05$, $**p < 0.01$, $***p < 0.001$, and $****p < 0.0001$.

**Acknowledgements**

This study was supported by funding from the NIH SARS-CoV-2 Assessment of Viral Evolution (SAVE) Program (subcontract no. 0258-A700-4609 under federal contract no. 75N93021C00014) to D.D.H. and (subcontract GR0010139-PO024016 under federal contract no. 75N93021C00016) to A.G., the Gates Foundation (project INV019355) to D.D.H., internal startup funding UR014016 from Columbia University to Y.G., K23 AI171263 to L.J.P., K24 AI155230 to M.T.Y., and K08 AI180347 to A.B.

We express our gratitude to Amanda Castillo, Meredith McNairy and Antonia Sturizo for conducting the C-PIC study (Columbia), and to Zijin Chu, Theresa Kowalski-Dobson, Anna Buswinka, Gabe Simjanovski, Joseph Wendzinski, Mayurika Patel, Kathleen Lindsey, and Dawson Davis of the VIVA study team for conducting the VIVA study.

**Author Contributions**

Experiments were conducted and data analyzed by Q.W., M.W., I.A.M., and Y.G. Serum samples were collected by I.A.M., A.B., J.G.S., L.J.P., C.G., R.V., M.T.Y., and A.G. The manuscript was written by Q.W., I.A.M., M.W., Y.G., and D.D.H., with feedback from all authors. All contributing authors have reviewed and endorsed the manuscript.

**Declaration of Interests**

D.D.H. co-founded TaiMed Biologics and RenBio, and he serves as a consultant for WuXi Biologics and Brii Biosciences and is a board director at Vicarious Surgical. A.G. served as a member of the scientific advisory board for Janssen Pharmaceuticals and has consulted and serves on a scientific advisory board for Sanofi Pasteur. The remaining authors declare no conflicts of interest.

**Supplemental Table and Figures**

**Table S1. Summary of clinical cohorts.**

No., number; WT, wildtype; MV, monovalent vaccine; BV, bivalent vaccine.

| | | KP.2 MV | |
|---|---|---|---|
| | | No. or Mean | % or (range) |
| **Total** | | 16 | |
| **Female** | | 9 | 56.3% |
| **Male** | | 7 | 43.8% |
| **Age** | | 41.4 | (21, 66) |
| **No. Vaccines** | All vaccines | 1.5 | (3, 8) |
| | WT | 3.0 | (2, 4) |
| | BA.5 BV | 0.8 | (0, 2) |
| | XBB.1.5 | 0.8 | (0, 2) |
| | KP. 2 MV | 1 | (1, 1) |
| **No. Infections** | | 1.5 | (0, 4) |
| **Days Post Infection** | | 679.6 | (195, 1352) |
| **Days Before KP.2 MV Vaccination** | | 14.4 | (0, 143) |
| **Days Post KP.2 MV Vaccination** | | 30.9 | (26, 39) |

**Table S2. Demographics of clinical cohorts.**

Vaccine formulations are denoted as wildtype (WT), BA.5 Bivalent (BA.5), XBB.1.5 monovalent (XBB.1.5), and KP.2 monovalent (KP.2). Vaccine manufacturers are denoted as Pfizer (P), Moderna (M), and Unknown (U). Yr, years; Infx, infection; Vax, vaccination; DBV, days before vaccination; DPV, days post vaccination; DPI, days post infection; F, female; M, male; Wh, white; Af, Black or African American; As, Asian.

| Participant ID | Age (Yr) | Sex | Race | DBV | DPV | No. Infx | DPI | No. Vax | | | | | Vaccine History |
|---|---|---|---|---|---|---|---|---|---|---|---|---|---|
| | | | | | | | | Total | WT MV | BA.5 BV | XBB.1.5 MV | KP.2 MV | |
| UMICH-1 | 35 | F | Wh | 1 | 29 | 2 | 1215 | 5 | 3 | 0 | 1 | 1 | WT-P/WT-P/WT-M/XBB.1.5-M/KP.2 MV-M |
| UMICH-2 | 29 | F | Af | 0 | 31 | 1 | 753 | 6 | 3 | 1 | 1 | 1 | WT-P/WT-P/WT-P/BA.5-P/XBB.1.5-M/KP.2-P |
| UMICH-3 | 39 | F | Wh | 7 | 30 | 1 | 984 | 6 | 3 | 1 | 1 | 1 | WT-P/WT-P/WT-P/BA.5-P/XBB.1.5-M/KP.2-P |
| UMICH-4 | 55 | F | Wh | 2 | 29 | 1 | 622 | 6 | 3 | 1 | 1 | 1 | WT-M/WT-M/WT-M/BA.5-M/XBB.1.5-M/KP.2-M |
| UMICH-5 | 58 | M | Wh | 10 | 31 | 2 | 407 | 7 | 4 | 1 | 1 | 1 | WT-P/WT-P/WT-P/WT-P/BA.5-P/XBB.1.5-P/KP.2-M |
| UMICH-6 | 37 | M | Wh | 3 | 28 | 0 | - | 5 | 3 | 1 | 0 | 1 | WT-M/WT-M/WT-P/BA.5-U/KP.2-U |
| CUMC-1 | 35 | M | As | 7 | 26 | 1 | 880 | 4 | 3 | 0 | 0 | 1 | WT-P/WT-P/WT-P/KP.2-M |
| CUMC-2 | 25 | F | As | 2 | 31 | 0 | - | 6 | 3 | 1 | 1 | 1 | WT-P/WT-P/WT-P/BA.5-M/XBB.1.5-M/KP.2-M |
| CUMC-3 | 21 | M | As | 0 | 37 | 4 | 195 | 3 | 3 | 0 | 0 | 1 | WT-P/WT-P/WT-P/KP.2-P |
| CUMC-4 | 29 | F | As | 18 | 28 | 1 | 244 | 4 | 3 | 0 | 0 | 1 | WT-P/WT-P/WT-P/KP.2-M |
| CUMC-5 | 55 | M | As | 1 | 30 | 2 | 451 | 5 | 3 | 1 | 0 | 1 | WT-M/WT-M/WT-M/BA.5-P/KP.2-P |
| CUMC-6 | 36 | M | Wh | 2 | 30 | 2 | 796 | 6 | 3 | 1 | 1 | 1 | WT-P/WT-P/WT-P/WT-P/BA.5-P/XBB.1.5-P/KP.2-P |
| CUMC-7 | 42 | M | As | 5 | 34 | 1 | 726 | 6 | 3 | 1 | 1 | 1 | WT-P/WT-P/WT-M/BA.5-M/XBB.1.5-M/KP.2-P |
| CUMC-8 | 40 | F | Wh | 30 | 28 | 3 | 240 | 6 | 3 | 1 | 1 | 1 | WT-M/WT-M/WT-M/BA.5-P/XBB.1.5-P/KP.2-M |
| CUMC-9 | 61 | F | Wh | 143 | 33 | 2 | 650 | 5 | 2 | 1 | 1 | 1 | WT-P/WT-P/BA.5-P/XBB.1.5-P/KP.2-M |
| CUMC-10 | 66 | F | Wh | 0 | 39 | 1 | 1352 | 8 | 3 | 2 | 2 | 1 | WT-P/WT-P/WT-M/BA.5-M/BA.5-M/XBB.1.5-M/XBB.1.5-M/KP.2-M |

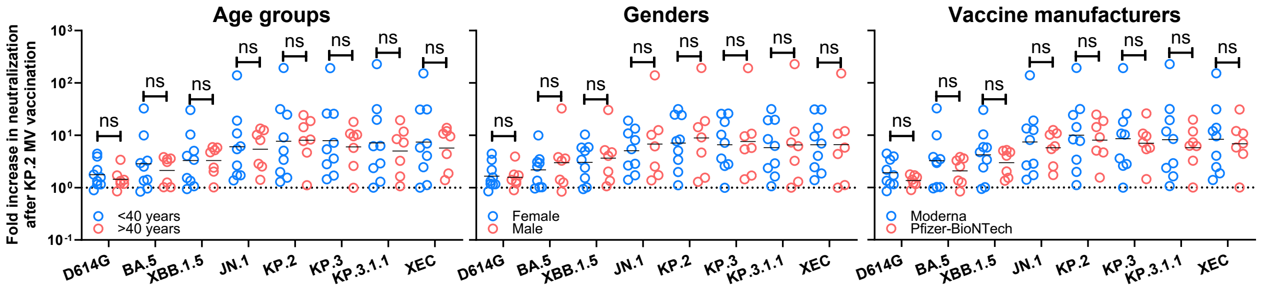

**Figure S1. Boost effects by KP.2 MV between age groups, self-reported genders, or vaccine manufacturers.**

Data are presented as fold changes in neutralization $ID_{50}$ titers following KP.2 MV vaccination for across age groups (<40 years vs. >40 years), self-reported genders (female vs. male), or vaccine manufacturers (Moderna vs. Pfizer-BioNTech). Bars represent geometric mean $ID_{50}$ titers and statistical analyses were performed using Mann-Whitney unpaired tests. ns, not significant.
